# Neural mechanism of experience-dependent sensory gain control in *C. elegans*

**DOI:** 10.1101/2021.12.14.472713

**Authors:** Yosuke Ikejiri, Yuki Tanimoto, Kosuke Fujita, Fumie Hiramatsu, Shuhei J. Yamazaki, Yuto Endo, Yasushi Iwatani, Koichi Fujimoto, Koutarou D. Kimura

## Abstract

Animals’ sensory systems adjust their responsiveness to environmental stimuli that vary greatly in their intensity. Here we report the neural mechanism of experience-dependent sensory adjustment, especially gain control, in the ASH nociceptive neurons in *Caenorhabditis elegans*. Using calcium imaging under gradual changes in stimulus intensity, we find that the ASH neurons of naive animals respond to concentration increases in a repulsive odor 2-nonanone regardless of the magnitude of the concentration increase. However, after preexposure to the odor, the ASH neurons exhibit significantly weak responses to a small gradual increase in odor concentration while their responses to a large gradual increase remain strong. Thus, preexposure changes the slope of stimulus–response relationships (*i*.*e*., gain control). Behavioral analysis suggests that this gain control contributes to the preexposure-dependent enhancement of odor avoidance behavior. Mathematical analysis reveals that the ASH response consists of fast and slow components, and that the fast component is specifically suppressed by preexposure. In addition, genetic analysis suggests that G protein signaling is required for the fast component. Thus, our integrative study demonstrates how prior experience dynamically modulates stimulus–response relationships in sensory neurons, eventually leading to adaptive modulation of behavior.

## INTRODUCTION

Animals use their sensory organs to interpret stimuli from the external environment and to effectively survive and reproduce. The intensity of these stimuli can vary by a factor of 10^10^, although the range of neuronal activity is generally limited to a factor of 10^2^ (Shapley and Enroth-Cugell, 1984). Thus, animals need to adjust their range of neuronal activity in peripheral and central sensory systems according to stimulus intensity. Such regulation of neuronal responsiveness has been reported in visual, auditory, olfactory, mechanical, and nociceptive systems, in a variety of animal species ranging from invertebrates to vertebrates (Carew et al., 1971; Dragoi et al., 2000; Priebe and Ferster, 2002; Root et al., 2008; Ulanovsky et al., 2003; Woolf and Ma, 2007).

One type of neuronal response modulation is sensory gain control. Gain control refers to a modulation that changes the slope of stimulus–response relationships and is different from adaptation or sensitization, which decreases or increases the overall responsiveness (Figure 1—figure supplement 1). Gain control has been reported to occur in the visual, auditory and somatosensory cortices of mammals and the olfactory circuit of *Drosophila*, and it is likely conserved across taxa (Andersen et al., 1985; Anderson et al., 2017; Azimi et al., 2020; Chance et al., 2002; Ohzawa et al., 1982; Olsen and Wilson, 2008). However, neuronal and/or molecular mechanisms underlying gain control, as well as the effect of gain control on sensory behavior, have not been sufficiently elucidated.

The nematode *Caenorhabditis elegans* has been widely used to study the mechanisms of sensory responses because of the feasibility in analyzing neural functions at the molecular, cellular, and behavioral levels. The animals respond to various sensory stimuli, such as chemicals, temperature, osmolality, and mechanical stimuli, which are modulated by experience as learning, and the neurons and genes involved in these responses have also been identified (Bargmann, 2006; De Bono and Maricq, 2005; Ferkey et al., 2021; Sasakura and Mori, 2013). However, most of the previous sensory experiments for physiological analyses of wild-type animals employed step-wise stimuli that were not controlled gradually in a physiologically meaningful manner. In addition, sensory gain control has been shown only in a few mutant strains (Kuhara et al., 2002; Saro et al., 2020). Thus, it has not been clear whether and how sensory gain control contributes to the wild-type animals’ sensory behavior, such as navigation under a chemical gradient.

*C. elegans* avoidance behavior to the odorant 2-nonanone is an ideal experimental paradigm to study the animals’ sensory response and experience-dependent modulation. The animals avoid the odor, and this odor avoidance behavior is enhanced by preexposure to the odor for 1 hour as a type of non-associative learning (Figure 1A) (Bargmann et al., 1993; Kimura et al., 2010). In previous studies, we found that the odor avoidance behavior consists of two behavioral states: (1) run, a long period of straight movement, and (2) pirouette, a period of repeated short movements with frequent directional changes (Kimura et al., 2010; Pierce-Shimomura et al., 1999). We also found that when the animals sense increases or decreases in odor concentration, it promotes pirouettes or runs, respectively (Figure 1—figure supplement 2A) (Tanimoto et al., 2017). In addition, calcium imaging of their neuronal activities with gradual changes in the odor concentration (Figure 1—figure supplement 2B) revealed that a pair of ASH nociceptive neurons and a pair of AWB olfactory neurons respond to the odor increase and decrease, respectively (Tanimoto et al., 2017). Furthermore, our pharmacological, genetic and mathematical analyses of neuronal activities have demonstrated that slowly increasing intracellular calcium response to a simple gradual odor decrease in AWB neurons is modeled by the leaky integration of time-differential of odor concentration (see Figure 1—figure supplement 2C; for leaky integration equation, please see Materials and Methods), which is mainly mediated by continuous influx of calcium ions through the L-type voltage-gated calcium channel (VGCC) EGL-19 (Tanimoto et al., 2017). Pharmacological analysis also suggested that the ASH response to a simple gradual odor increase (odor gradient #1) consists of an AWB-like slow component plus a fast and transient component, which has not been modeled yet (Figure 1—figure supplement 2D) (Tanimoto et al., 2017). We further found that the ASH response to a small odor increase is reduced by odor preexposure, indicating an experience-dependent modulation of its activity (Yamazaki et al., 2019). ASH neurons are considered as a simple model for polymodal nociceptive neurons evolutionarily conserved from worms to mammals (Kaplan and Horvitz, 1993). Thus, analyzing the ASH response to 2-nonanone will be ideal to address questions at the levels of behavior, neural activity, and molecules. For example, how do sensory neurons adapt their sensitivity for behavioral responses to ever-changing environmental stimuli, which cellular mechanisms support changes in response after experiencing a stimulus, and what is the molecular mechanism for its implementation.

**Figure 1.**
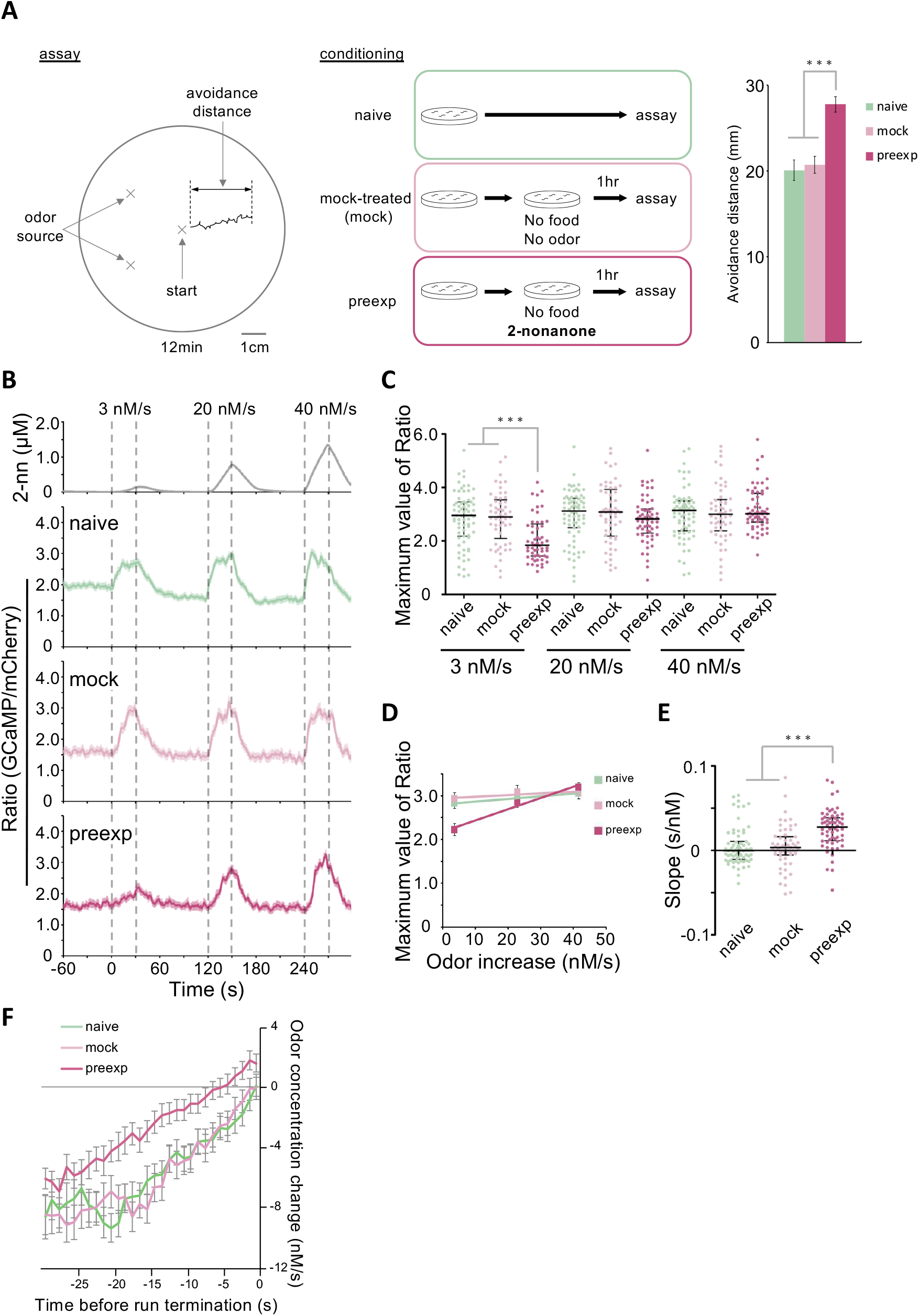
Preexposure to the repulsive odor 2-nonanone causes the enhancement of 2-nonanone avoidance and sensory gain control in ASH neurons.**A**, Preexposure-dependent enhancement of odor avoidance behavior. (left) Example of an animal’s trajectory tracked during 12 minutes of 2-nonanone avoidance assay. (middle) The three conditions for 2-nonanone avoidance assay. (right) Result of 2-nonanone avoidance assay, where the average avoidance distance ± standard error in each condition is shown (n = 47, 45, 44 in naive, mock-treated, and preexposed conditions, respectively). Compared to control (*i*.*e*., naive and mock-treated) animals, preexposed animals exhibit significantly longer avoidance distances. *** *p* < 0.001 (Kruskal-Wallis test with *post hoc* Steel-Dwass test). **B**, ASH responses to three consecutive stimuli (3, 20, and 40 nM/s; top) in the three conditions. In the naive (green) and mock-treated (pale red) conditions, the responses were always large regardless of the stimulus intensity. However, in the preexposure condition (dark red), the responses were smaller for small stimuli and larger for large stimuli. The solid lines and associated shadows are the average values and their standard errors, respectively (n = 71, 57, 61 in naive, mock-treated, and preexposed conditions, respectively). The black dotted lines indicate the onset and end of the odor increase phase. **C**, Scatter plot of the maximum values of ASH response (shown in panel B) to each stimulus. *** *p* < 0.001 (Kruskal-Wallis test with *post hoc* Steel-Dwass test). **D**, Correlation between the rate of odor concentration increase and the maximum value of ASH response. The mean ± standard error of ASH maximum values in panel C and its linear approximation are shown. **E**, Scatter plot of the slope of the response. Each slope was calculated by linear approximation of maximum values of ASH responses of an animal to the three stimuli (3, 20, and 40 nM/s). *** *p* < 0.001 (Kruskal-Wallis test with *post hoc* Steel-Dwass test). **F**, Time-course changes of odor concentration that the animals sensed before the initiation of a pirouette. The odor concentrations that each animal sensed 30 seconds before the initiation were ensemble-averaged. The average values ± standard error of naive, mock-treated, and preexposed animals are shown in green, pale red and dark red, respectively (n = 50 in all conditions). In the naive and mock-treated animals, the transition from run to pirouette on average occurred when the average value of odor concentration change became close to zero, whereas the transition occurred several seconds after the animals started sensing increases in the odor concentration in the preexposed condition. The statistical details are described in Supplemental Table 1. **Figure 1—figure supplement 1:** Schematic drawing of sensory modulations. **Figure 1—figure supplement 2:** Behavioral and neuronal responses of *C. elegans* to the repulsive odor 2-nonanone. **Figure 1—figure supplement 3:** Cartoons of experience-dependent modulation of the odor avoidance behavior.

In this study, we show that sensory gain control occurs in ASH sensory neurons of *C. elegans* for efficient odor avoidance by preexposure to 2-nonanone via specific suppression of one of its activity components. Subjecting the animals to a series of gradual changes in odor concentration reveals that preexposed ASH neurons become less sensitive to smaller increases in odor concentration, while their responses to larger increases do not change. In other words, preexposure changes the slope of the stimulus–response relationships, i.e. gain control. This experience-dependent modulation of sensory activity is consistent with the changes in the animals’ behavioral responses: they respond less to small increases in odor concentration, likely leading to more efficient odor avoidance not disturbed by a noisy and/or fluctuating odor concentration change. On modeling the ASH response mathematically, we find that the slow and fast components of ASH response are well fitted as the sum of the leaky integration of the first- and second-order time-differentials of odor concentration, respectively. Notably, the preexposure experience specifically suppresses only the fast component. Furthermore, genetic analysis suggests that G protein signaling may regulate the fast component after preexposure. Thus, our results demonstrate how experience-dependent modulation of stimulus–response relationships occurs and how it leads to changes in behavioral responses.

## RESULTS

### Sensory gain control in ASH neurons caused by odor preexposure

To reveal how experience changes the activities of sensory neurons under conditions of physiologically meaningful odor stimuli, we investigated ASH activities before and after preexposure to 2-nonanone. Our previous quantitative behavioral analysis revealed that *C. elegans* senses approximately 5–20 nM/s concentration changes during 2-nonanone avoidance behavior (Yamazoe-Umemoto et al., 2015). Therefore, in this experiment, we used three types of stimuli in a series of odor stimulations (odor gradient #2): (1) a minimum increase that could be provided stably by our original microscope system OSB2 (3 nM/s) (Tanimoto et al., 2017); (2) a maximum increase sensed during the behavioral experiment (20 nM/s); and (3) an even larger increase (40 nM/s). We measured the ASH response to these stimuli under three different conditions with prior treatments: naive, mock-treated, and preexposed (Figure 1A).

In the naive condition, the ASH response to 3 nM/s started to increase immediately after the onset of odor increase, quickly reached close to the maximum value, and the magnitude of the response was sustained during the odor increase (Figure 1B). When the odor began to decrease, the response also decreased, and when the odor concentration returned to zero, the ASH response returned to a basal level. The ASH responses to 20 nM/s and 40 nM/s odor increases were essentially similar to the response to 3 nM/s in magnitude and pattern. The results of the mock-treated condition were similar to those of the naive animals. In summary, the magnitude of the response was always constant in naive and mock-treated conditions, regardless of the magnitude of the odor increase.

In the preexposed condition, unlike naive and mock-treated animals, ASH did not respond much to the 3 nM/s increase: its maximum value was substantially smaller than those in the naive and the mock-treated animals, and the activity slowly increased and reached the maximum value at the end of the odor-increasing phase (Figure 1B). However, ASH responded strongly to the 20 and 40 nM/s increases, and the magnitude of the response increased according to the magnitude of the odor increase.

In terms of the maximum response to each odor increase, there was no significant difference in the responses to 20 and 40 nM/s, but the response to 3 nM/s was significantly smaller in the preexposed animals than in the naive and mock-treated animals (Figure 1C). The linear approximation of the average responses to each odor increase did not change much in the naive and mock-treated conditions, but was positive in the preexposed condition (Figure 1D). Furthermore, by calculating the slope of each animal’s response to the 3, 20, and 40 nM/s stimuli, we found that the slopes of the preexposed response were significantly larger than those of the naive and mock-treated responses (Figure 1E). Thus, the slope of stimulus– response relationships of ASH neurons changes because of preexposure to the odor, not adaptation or sensitization, which is an overall increase or decrease of the relationships without changes in its slope (Figure 1—figure supplement 1). In other words, gain control occurs in ASH neurons because of preexposure to the odor.

### Behavioral significance of ASH gain control by preexposure

Next, we investigated whether the ASH gain control after preexposure could explain the enhanced 2-nonanone avoidance behavior (Kimura et al., 2010). Since 2-nonanone is a repulsive odor and ASH responds to the odor increase to cause turns (Tanimoto et al., 2017), a simple scenario would be that ASH sensitivity increases to make the animal avoid 2-nonanone sooner after preexposure. However, our results indicate that ASH was less sensitive when the increase in odor concentration was small (Figure 1B).

To understand how changes in ASH response caused by gain control affect odor avoidance behavior, we calculated time-course changes in odor concentration that each animal sensed during the odor avoidance. This calculation was according to the model of 2-nonanone evaporation and diffusion, which is based on the actual measurement of the local odor concentration in the air phase of the plate (Tanimoto et al., 2017). We then analyzed the correlations between odor concentration changes and behavioral aspects of the animals.

We found that the change in odor concentration upon initiation of the pirouette phase was almost zero in the naive and mock-treated conditions, but was positive in the preexposed condition (Figure 1F). This suggests that naive and mock-treated animals initiated pirouettes in response to very small increases in odor concentration, while the preexposed animals responded only to larger increases. This is consistent with our results on the ASH response, which is very sensitive to a small odor increase in naive and mock-treated conditions, but less sensitive in the preexposed condition (Figure 1B–E). This change can explain the enhanced odor avoidance behavior by the animals after odor preexposure (Figure 1—figure supplement 3) (see Discussion).

### Modeling the experience-dependent gain control in ASH activity

In order to obtain quantitative insights into the mechanism of gain control caused by preexposure, we performed mathematical modeling of the ASH response. In our previous study, the ASH response to the simple gradual odor increase (odor gradient #1) was modeled by simple time-differential of odor concentration (Tanimoto et al., 2017). Furthermore, our pharmacological study suggested the ASH response to the odor gradient #1 consists of fast and slow components (Figure 1—figure supplement 2D). The slow component was modeled by the leaky integration of time-differential (*dC/dt*) of odor concentration (*C*) like the AWB response (Figure 1—figure supplement 2C). However, the simple time-differential model and the leaky integration of time-differential model (Equation 1 in Materials and Methods) did not reproduce the ASH responses sufficiently, especially the fast component of the naive and mock-treated ASH responses to the odor gradient #2 (Figure 2—figure supplement 1 and 2).

Next, we sought to model the fast component of ASH response mathematically. The fast component started to increase at the onset of odor concentration increase, and soon it started to decrease even when the odor concentration continued to increase (red dotted line in Figure 1—figure supplement 2D). This time course of the fast component can be approximated by a leaky integration of the second-order time-differential (*d*^*2*^*C/dt*^*2*^) of the odor concentration (vertical gray bar and red line in “*d*^*2*^*C/dt*^*2*^” panel in Figure 2A). To this end, we extended the model, hereafter referred to as “first and second differential model”, where the ASH response itself is represented by a leaky integration of the sum of both the first- and second-order time-differentials of the odor concentration (Figure 2A; Equation 2 in Materials and Methods).

**Figure 2.**
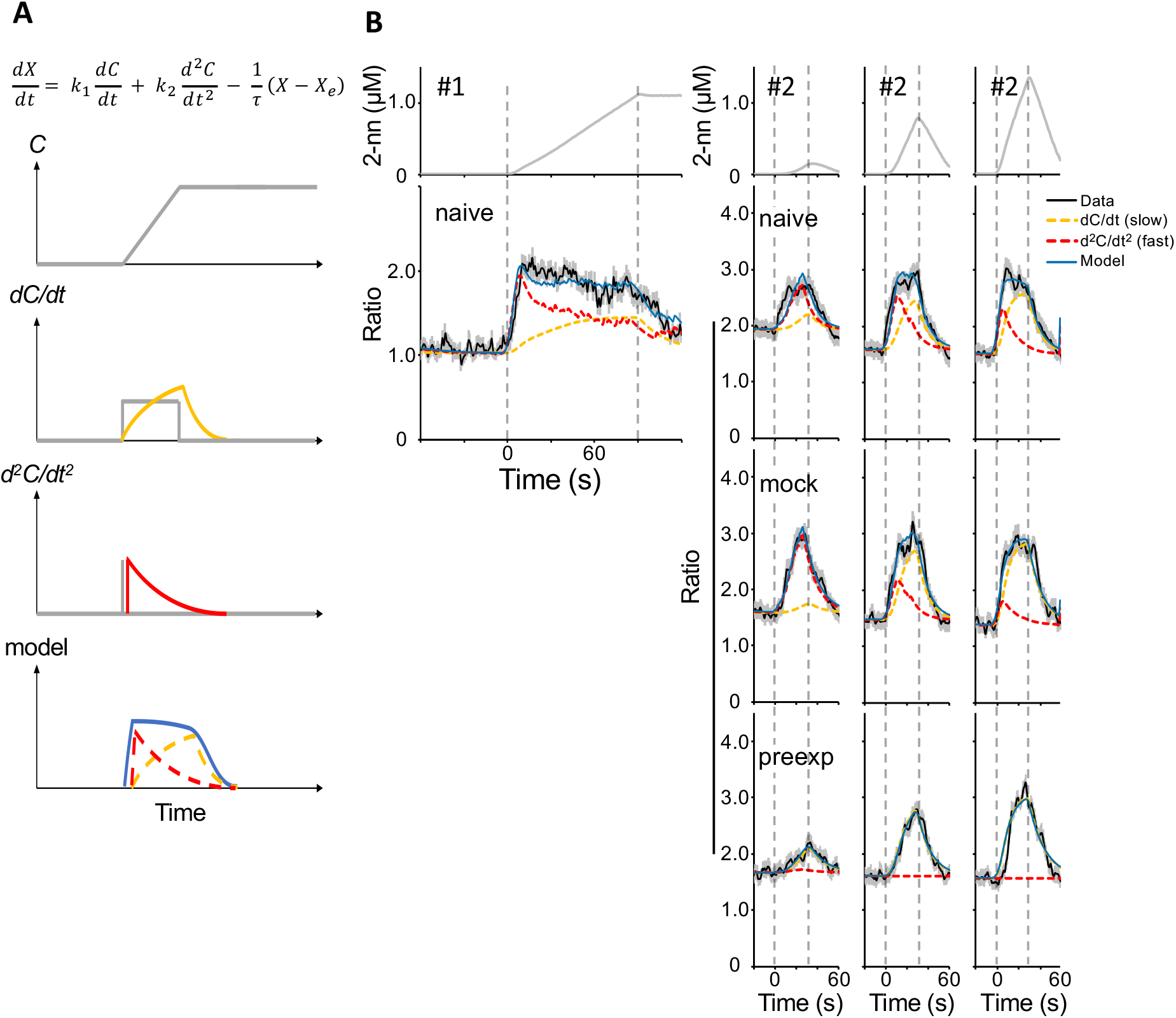
Mathematical model of wild-type ASH response independently fitted to a single odor stimulus.**A**, Schematic drawing of a mathematical model of ASH response. Equation of the first and second differential model is shown at the top. Also shown are the time course of a simple gradual odor concentration increase (top panel), first-order time-differential of odor concentration and its leaky integration (gray rectangle and yellow line, respectively, in the second panel), second-order time-differential of the odor concentration and its leaky integration (gray vertical bar and red line, respectively, in the third panel; negative value of second-order time-differential is not calculated), and the sum of leaky integration of first- and second-order time-differentials (blue solid, yellow dashed and red dashed lines, respectively, in the bottom panel). **B**, The results of fitting the ASH response to the simple gradual odor increase used in the previous study (left; odor gradient #1, n = 39) and independent fitting to each of three consecutive odor stimuli used in Figure 1B (right; odor gradient #2, n = 71, 57, 61 in naive, mock-treated, and preexposed conditions, respectively). The top panels exhibit each odor concentration change, and lower panels exhibit ASH responses of different conditions of animals (naive, mock-treated or preexposed) to each stimulus. The black line and the associated gray area are the average ASH responses and their standard errors, the blue line is the model, the yellow and red dashed lines are the first- and second-order components in the model, respectively. The black dotted line indicates the onset and end of the odor increase phase. **Figure 2—figure supplement 1:** Independent fitting of the wild-type ASH response to the odor stimuli with the original simple time-differential model. **Figure 2—figure supplement 2:** Independent fitting of the wild-type ASH response to the odor stimuli with the leaky integration of first-order time-differential model. **Figure 2—figure supplement 3:** Independent fitting of the wild-type ASH response to the odor stimuli with the leaky integration of second-order time-differential model.

We examined whether the first and second differential model could reproduce the ASH responses to odor gradients #1 and #2. By determining the model parameters to the odor gradient #1 and each odor stimulus in odor gradient #2 independently, in most of the conditions the model reproduced the ASH activities well, including the fast component (blue lines in Figure 2B). We should note that the contribution of the fast component in ASH activities was significantly lower in preexposed than in naive and mock-treated conditions (red lines in right panels in Figure 2B), indicating that the fast component was specifically suppressed by the preexposure (see below).

To test the optimality of the first and second differential model, we compared the goodness of fit among the first and second differential model and three other models: the original simple time-differential model, and leaky integration of only first- or second-order time-differential models (Figure 2—figure supplement 1, 2 and 3, respectively). By calculating the Bayesian Information Criterion (BIC) (Schwarz, 1978) for each model, we found that the first and second differential model had the best or close-to-the-best goodness of fit in all naive and mock-treated conditions (Table 1), demonstrating the optimality of the model for these conditions. In the preexposed condition, the leaky integration of first-order time-differential model had the best fit, consistently with the suppression of the fast (second-order time-differential) component in ASH activity (Figure 2B right).

**Table 1.** Summary of comparing BIC value among mathematical models.

| Odor | #1 | 3nM/s increase in #2 |  |  | 20nM/s increase in #2 |  |  | 40nM/s increase in #2 |  |  |
| --- | --- | --- | --- | --- | --- | --- | --- | --- | --- | --- |
| Condition | Naive | Naive | Mock-treated | Preexp | Naive | Mock-treated | Preexp | Naive | Mock-treated | Preexp |
| $X = k \frac{dC}{dt} + X_e$ | -557.3 | -216.9 | -195.0 | -278.5 | -153.5 | -125.8 | -140.8 | -158.9 | -92.5 | -68.8 |
| $\frac{dX}{dt} = k_1 \frac{dC}{dt} - \tau(X - X_e)$ | -559.9 | -240.4 | -217.7 | **-423.4** | -205.0 | -233.8 | **-379.2** | -235.6 | -260.8 | **-261.5** |
| $\frac{dX}{dt} = k_2 \frac{d^2C}{dt^2} - \tau(X - X_e)$ | -741.1 | **-287.6** | **-368.6** | -373.8 | -253.5 | -211.6 | -211.3 | -224.4 | -159.4 | -109.7 |
| $\frac{dX}{dt} = k_1 \frac{dC}{dt} + k_2 \frac{d^2C}{dt^2} - \tau(X - X_e)$ | **-769.6** | -286.5 | -366.7 | -420.5 | **-301.2** | **-298.1** | -374.8 | **-336.3** | **-289.0** | -257.1 |

BIC values were calculated independently for each odor stimulus. The conditions with the smallest BIC values under each condition are
indicated in bold.

The contribution of the first and second differential exhibited a remarkable difference for every odor gradient #2 (Figure 2B right). For example, in naive and mock-treated conditions, the fast component became smaller when odor concentration increased more rapidly (compare the response to the first odor stimulus to the ones to the second and third odor stimuli). Consistently, when these parameters were uniquely determined for all the odor stimuli, the model did not accurately reproduce the ASH responses to odor gradient #2 (Figure 3—figure supplement 1). These results suggest that the parameters depend on some aspects of the stimuli such as the odor concentration and/or its changing velocity.

To test whether and how the parameters depend on aspects of stimuli, we plotted the relationships between the parameters and multiple aspects of the odor stimuli. We found that each parameter approximately follows a logarithmic function of the stimulus intensity (Figure 3—figure supplement 2). Interestingly, in the well-fitted cases (red rectangles in Figure 3— figure supplement 2), this function of the slow component (*k*_*1*_) and the leaky part (1/*τ*) were similar among naive, mock-treated and preexposed conditions, although that of the fast component (*k*_*2*_) was much lower for preexposed than for naive and mock-treated conditions. When we further introduced this stimulus intensity dependency into the first and second differential model (Equation 3 in Materials and Methods), the model unifyingly reproduced the ASH responses to odor gradients #1 and #2 as well as their gain control-dependent changes (Figure 3A and B). These results suggest that the contributions of fast and slow components and the leak of ASH response depend on the logarithmic function of the odor stimulus intensity and that the fast component is further suppressed after odor preexposure for the gain control. The differences in the relationships between stimuli aspects and each parameter may reflect the regulatory mechanisms of fast and slow components and the leak activity (see Discussion).

**Figure 3.**
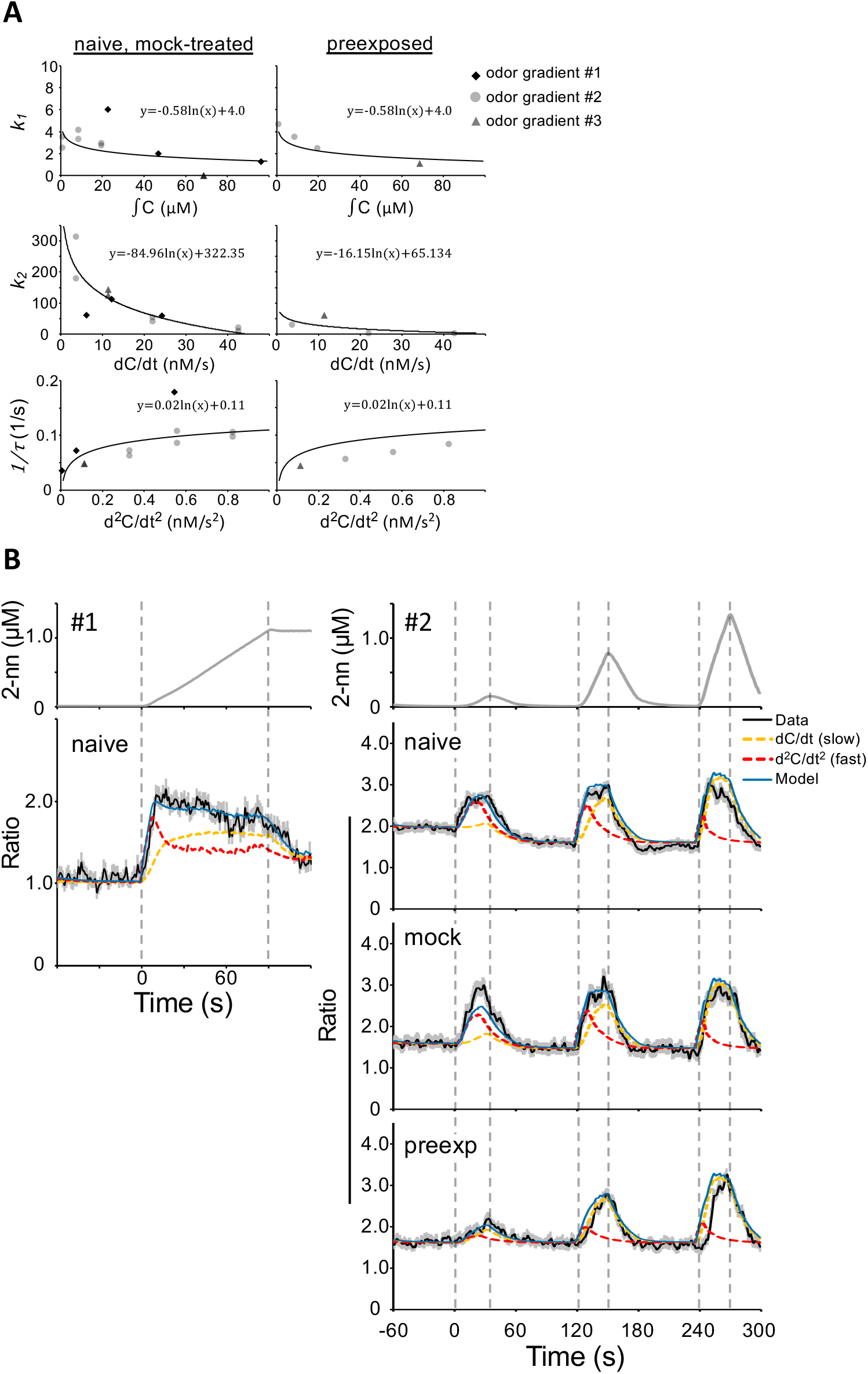
Model of wild-type ASH response fitted to the odor gradient #1 and #2 with stimulus-dependent parameters.**A**, Scatter plots used for the first and second differential model with stimulus-dependent parameters. Only the parameter of the second-order time-differential, *k*_*2*_, is substantially different between naive, and mock-treated versus preexposed conditions. Black rhombuses, light gray circles, and dark gray triangles represent each ASH response to odor gradient #1, #2, and #3, respectively. Odor gradient #3 was used for the genetic analysis (Figure 4). **B**, ASH response and its model to a simple odor increase (left ; odor gradient #1, n = 39) and to the three consecutive stimuli (right; odor gradient #2, n = 71, 57, 61 in naive, mock-treated, and preexposed conditions, respectively). The black line and the associated gray area are the average ASH responses and their standard errors, the blue line is the model, the yellow and red dashed lines are the first-and second-order component in the model, respectively. The black dotted line indicates the onset and end of the odor increase phase. **Figure 3—figure supplement 1:** Fitting of the model with constant coefficients for whole naive ASH response to the odor gradient #2. **Figure 3—figure supplement 2:** Scatter plots of the relationship between each input and parameter.

### Genetic analysis of the ASH response

Finally, we tried to identify the genes responsible for the fast component of ASH responses. Specifically we looked for mutants that exhibit the slow component-like response in naive or mock-treated (*i*.*e*., control) conditions.

We first analyzed mutants of cation channels. *osm-9* and *ocr-2* are the homologs of TRPV cation channels and known to function in ASH neurons for depolarization caused by sensory stimuli (Colbert et al., 1997; Tobin et al., 2002) although their physiological role in 2-nonanone sensation has not been revealed. *osm-9 ocr-2* double mutants exhibited substantially no response (Figure 4A), suggesting that they are also required for sensory depolarization caused by 2-nonanone. The L-type VGCC subunit α1 EGL-19 is responsible for the slow component, although loss-of-function mutations in N- and T-types of VGCC subunit α1, *unc-2* and *cca-1* respectively, did not affect ASH response to the odor (Tanimoto et al., 2017). In this study we tested the homologs of a VGCC auxiliary subunit α2δ2 and a sodium leak channel, *tag-180* and *unc-77*, respectively (Lainé et al., 2011; Yeh et al., 2008). Both of the mutant strains exhibited wild-type like responses that were properly fitted with the ASH model (Figure 4A), indicating that these genes are not involved in the ASH response either.

**Figure 4.**
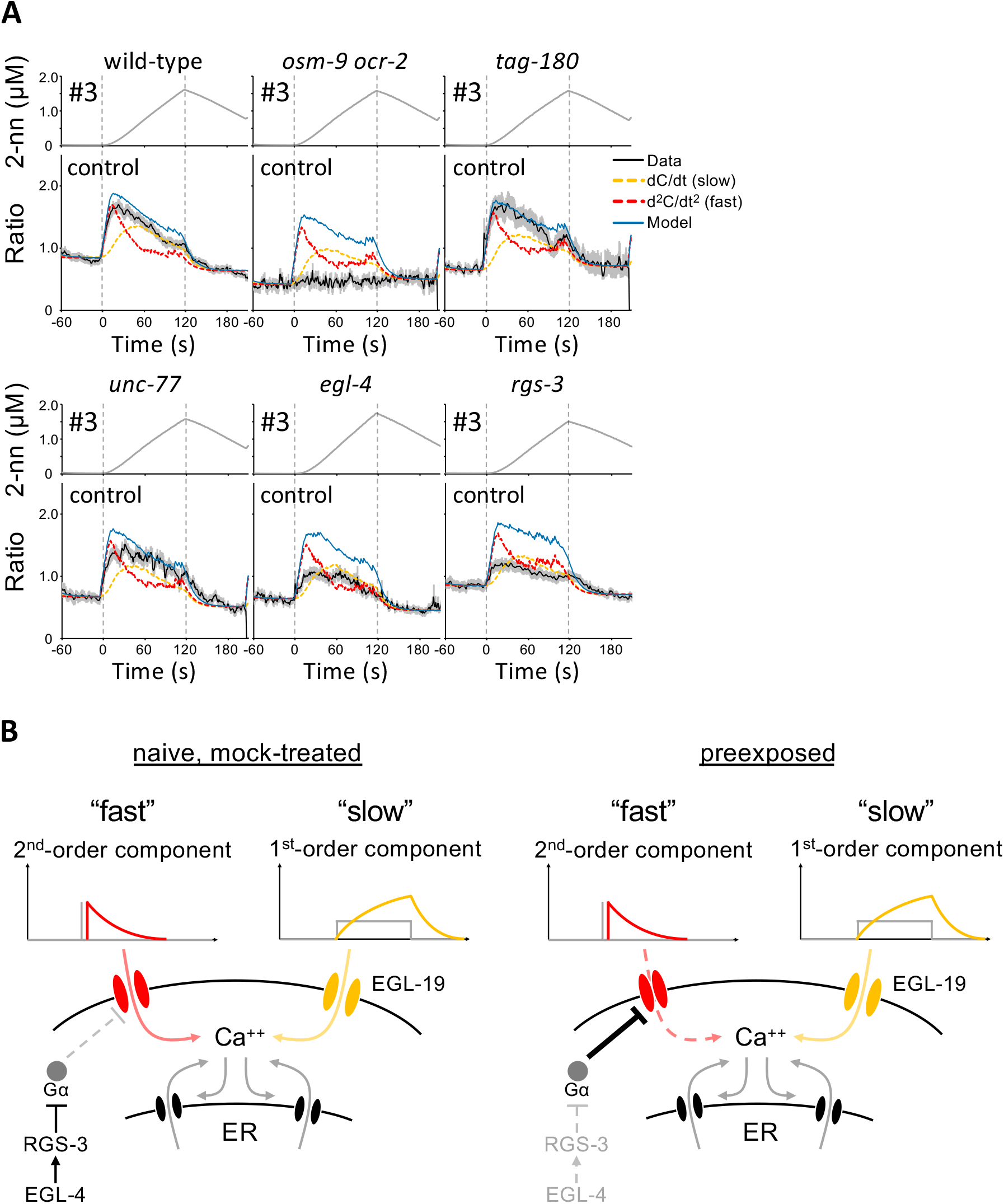
Genetic analysis of the ASH gain control. **A**, ASH response in wild-type and mutant animals in control condition (n =51 in wild-type; n = 7 in *osm-9 ocr-2*; n = 15 in *tag-180*; n = 16 in *unc-77*; n = 46 in *egl-4*; n = 29 in *rgs-3*) to a longer and larger continuous odor increase (up to ∼2 µM within 120 s; odor gradient #3). We used this odor stimulus with the idea that even milder mutant phenotypes would be observed clearly with a longer and larger stimulus. The black line and the associated gray area are the average ASH responses and their standard errors. The model parameters to fit for the wild-type response in Figure 3 were used, and overlayed with each of the ASH responses. The black dotted line indicates the onset and end of the odor increase phase. **B**, A proposed molecular model of ASH response. ASH response is mainly mediated by transiently active calcium channels (red) and persistently active L-type VGCCs (yellow). Gα protein activity can suppress the transiently active calcium channels, and this Gα protein activity is inhibited by PKG–RGS pathway in naive/mock-treated conditions. Gain control is caused only by suppression of the transiently active calcium channels via the Gα signaling. The calcium influx from the plasma membrane is further increased by the calcium-induced calcium release through IP3 receptor and ryanodine receptor on ER membrane (Tanimoto et al., 2017).

We then analyzed other candidates that have been known to be involved in modulation of sensory neuronal activity in *C. elegans*: *egl-4* (cyclic GMP-dependent protein kinase: PKG), and *rgs-3* (regulator of G protein signaling: RGS) (Ferkey et al., 2007; L’Etoile et al., 2002). The ASH responses of *egl-4* and *rgs-3* mutants in the control condition were substantially suppressed (black lines in Figure 4A). Remarkably, the ASH responses of *egl-4* and *rgs-3* mutants were best fitted with the first-order (*i*.*e*., slow) only model (yellow lines in Figure 4A and Table 2), suggesting that the fast component of ASH activity requires these gene products. It has been reported that EGL-4 phosphorylates RGS-3, and loss-of-function mutations in either of the genes increases ASH neuronal and ASH-mediated behavioral responses to other repulsive stimuli (Krzyzanowski et al., 2013). Therefore, this PKG–RGS pathway may regulate the fast component of the ASH response in a different manner from this previous report (see Discussion).

**Table 2.**
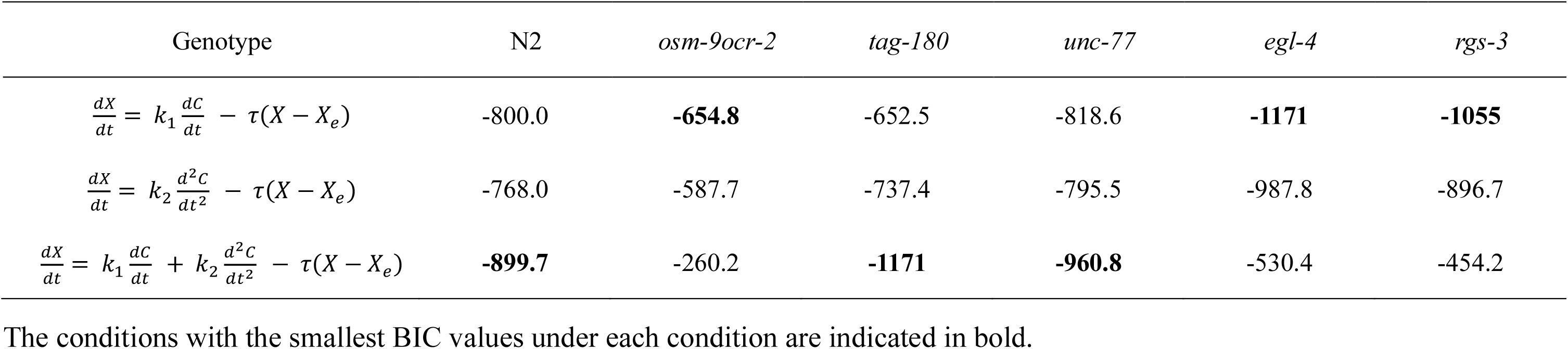
Summary of BIC value for the ASH mathematical model in mutants.

## DISCUSSION

### 2-nonanone avoidance enhancement by gain control in ASH sensory neurons

In this study, we demonstrated that the responsiveness of the ASH sensory neurons of *C. elegans* to 2-nonanone exhibits gain control after preexposure to the odor. In naive and mock-treated conditions, ASH always exhibits a large response regardless of the rate of odor increase, whereas in the preexposed condition, the response to a small gradual increase became smaller and the response to a large gradual increase remains unchanged (Figure 1B– E). This suggests that gain control to change the slope of stimulus–response relationships is caused by experiencing odor preexposure.

Based on previous findings (Tanimoto et al., 2017) and the results of this study (Figure 1F), we consider that this sensory gain control contributes to the ability of *C. elegans* to efficiently navigate down the gradient of repulsive odor. The animals in the 2-nonanone avoidance assay generally move away from the odor source and sense decreases in the odor concentration. However, their movements away from the odor source are frequently interrupted because of stochastic occurrence of pirouettes and/or of small increases in odor concentration (Yamazoe-Umemoto et al., 2015). This increase of odor concentration could be caused by (1) the direction of the animals’ movement straying from the ideal, (2) small fluctuations in the odor gradient, and/or (3) small periodic changes in the sensed odor concentration caused by the animals’ sinusoidal head swing. In the naive and mock-treated conditions, the animals initiate pirouettes when they experience even a small increase in odor concentration (Figure 1— figure supplement 3, top left). However, in the preexposed condition, the animals do not respond to small increases (Figure 1—figure supplement 3, top right), which results in a longer avoidance distance traveled in the same period. They still initiate pirouettes when they sense larger increases (Figure 1—figure supplement 3, bottom panels). Thus, experience-dependent changes in ASH response due to gain control likely contribute the enhanced odor avoidance behavior, which could also be effective in a real-world environment infused with noise.

To our knowledge, this is the first report of sensory gain control in wild-type *C. elegans*. There are multiple examples of behavioral changes in *C. elegans* due to activity changes in sensory neurons that are affected by repeated stimuli, different types of stimuli, or the feeding state (Chew et al., 2018; Ezcurra et al., 2016; Larsch et al., 2015). However, most of the response changes in wild-type animals are caused by adaptation or sensitization but not gain control. It is possible that the animals might have developed a special mechanism against this odorant to effectively avoid danger because it is one of the two major volatile compounds produced by *P. aeruginosa*, a bacterial species that is pathogenic to *C. elegans* (Labows et al., 1980; Tan et al., 1999; Tan and Ausubel, 2000).

### Estimating the neural mechanism of gain control by mathematical modeling

To obtain quantitative insights into the mechanism of sensory gain control, we performed mathematical modeling. Indeed, mathematical modeling has successfully revealed essential features of the sensory neuronal responses of the animals (Ikeda et al., 2021; Itskovits et al., 2018; Kato et al., 2014; Tanimoto et al., 2017). Our previous pharmacological analysis revealed that ASH activity is composed of fast and slow components, and we showed in this study that the fast component can be approximated by a leaky integration of the second-order time-differential of the odor concentration (Figure 2). Even independent of the pharmacological result, the ASH responses in wild-type animals exhibit overshoot in response to a long-lasting and linearly increasing odor concentration (odor gradient #1 in Figure 1— figure supplement 2D and Figure 2B, and odor gradient #3 in Figure 4A). More precisely, the response first increases rapidly, and then slowly decreases to a non-zero constant value. The leaky integration of the first-order time-differential of the odor concentration generates a sustained response that converges to a non-zero constant value. The leaky integration of the second-order time-differential generates a rapidly increasing and subsequently decreasing response. Thus, both the pharmacological analysis as well as the analysis of shapes of ASH response in time domain suggest that the ASH response can be modeled with a leaky integration of the sum of the first- and second-order time-differentials of the odor concentration, and that these two terms well describe essential characteristics of ASH neurons. Indeed, the model had the best goodness of fit in most naive and mock-treated conditions in terms of BIC (Table 1). It should be noted, however, that we only tested the leaky integration of the second-order time-differential equation to the constantly increasing odor gradients, and the equation may not be applicable to ASH responses to other types of odor stimuli; in other words, the fast component could be approximated with another equation such as an alpha function (Rall, 1967). Nevertheless, the leaky integration of the sum of the first- and second-order time-differential equation with stimulus-dependent parameters nicely reproduced the ASH responses, and provided us an important biological insight (see next section). Some details still do not match, suggesting the involvement of other factors in the ASH response.

Remarkably, our model indicates that the preexposure-dependent changes in ASH response can be caused only by the modulation of the fast component (*k*_*2*_) rather than the modulation of overall ASH response, although the ASH response does exhibit changes in multiple aspects, such as its speed, magnitude, and small increase/decrease when it reaches closes to the maximum value (odor gradient #2 in Figure 2B). It also explains why ASH responses to a large odor increase are not substantially affected. This result suggests that a specific sensory signaling molecular pathway that suppresses the fast component in ASH neurons is regulated by preexposure to cause experience-dependent modulation of behavior, *i*.*e*., learning.

### Relationships between the mathematical model and molecular mechanisms of ASH response

We have previously shown that AWB responses are mediated by EGL-19, an α1 subunit of L-type VGCC, which possesses a long-lasting channel opening property, and that a Gα_o_ homolog, GOA-1, is required for the first-order time-differential of odor concentration. These results suggest that the constant decrease in 2-nonanone concentration is time-differentiated at the sensory ending by G protein signaling and causes constant depolarization, leading to EGL-19 opening and a continuous influx of calcium into the cells. The slow component in ASH neurons is also mediated by EGL-19 (Tanimoto et al., 2017).

In contrast, the fast component in ASH neurons is likely caused by a transient calcium influx at the onset of odor concentration increase, which ends rapidly (Figures 2A and 4B left). One candidate gene product for this was the T-type VGCC (“T” for transient opening) (Nowycky et al., 1985). However, the loss-of-function mutant of the *cca-1* gene, the sole homolog of T-type VGCC in *C. elegans* (Steger et al., 2005), showed wild-type-like ASH responses (Tanimoto et al., 2017). In addition, the loss-of-function mutants of N-type homolog *unc-2* (Tanimoto et al., 2017) and of the homologs of VGCC auxiliary subunit α2δ2 *tag-180* (Figure 4A) also exhibited a wild-type-like response, suggesting that some other calcium channel(s) may be involved in the fast component.

Still, we found that PKG EGL-4 and RGS RGS-3 possibly modulate the fast component. RGS is known to inhibit G protein signaling to modulate the magnitude and/or time-course of neuronal responses in *C. elegans* and mammals (Cao et al., 2012; Krzyzanowski et al., 2013; Lur and Higley, 2015). In addition, PKG phosphorylates RGS to increase its activity in *C. elegans* and mammals (Huang et al., 2007; Krzyzanowski et al., 2013). Because the effects of PKG–RGS activation on neuronal responses are different between the previous reports and our result, we consider that these gene products may inhibit G protein activity which suppresses the ASH response. In other words, in naive/mock-treated ASH neurons, the PKG– RGS pathway inhibits G protein activity that suppresses the ion channel that mediates the fast component, and preexposure suppresses the PKG–RGS pathway, leading to activation of G protein and to inhibition of the fast component (Figure 4B).

While genetic analysis itself is static (*i*.*e*., it would not provide us information related to time-course changes of the gene product activities), the combination with physiological analysis and mathematical modeling provides insights as to how these gene products affect neural activities dynamically. For example, the characteristics of ASH neurons in wild-type animals are not clear except for the preexposure-dependent gain control aspect, although a simple “first and second differential model” can explain multiple aspects of the differences, such as the quick rise at the onset of the odor stimulus, the gradual decrease during a long-lasting odor increase, the slow decay during odor decrease (not reproduced by the original simple time-differential model) as well as the gain control (Figures 2 and 3B). Furthermore, the stimulus-dependent parameters we used in the model may reflect the opening probabilities of the channels for the fast and slow responses, which change according to the intensity of the stimulus. For example, *k*_*1*_ reduces according to the accumulation of odor concentration (Figure 3A), which may indicate habituation due to the sustained stimulation from the odor. Further, *k*_*2*_ reduces according to the average of *dC/dt*, suggesting that the depolarization level may affect the opening probability because the odor concentration is time-differentiated at the sensory ending and it reflects the depolarization level (Tanimoto et al., 2017). Lastly, 1/*τ* increases according to the second-order time-differential of odor concentration, which may reflect a change in leakage level at the stimulus onset. Those insights could have not been obtained without the model.

In summary, our results demonstrate that gain control occurs in ASH sensory neurons after preexposure to the repulsive odor, and mathematical analysis suggests that this gain control is caused by suppression of the fast component, which could be caused by transient cation influx at the onset of odor concentration increase. This gain control leads to efficient avoidance behavior that allows the animal to ignore slight increases in odor concentration while moving down the odor gradient, thereby enhancing the avoidance distance. In more complex animals, such as mice and *Drosophila*, neurotransmitters such as serotonin or GABA are involved in gain control in peripheral and central sensory systems (Azimi et al., 2020; Olsen and Wilson, 2008) but little is known about their detailed molecular mechanism. This study may contribute to our understanding of intracellular mechanisms surrounding sensory gain control in animals.

## MATERIALS AND METHODS

### Key Resources Table

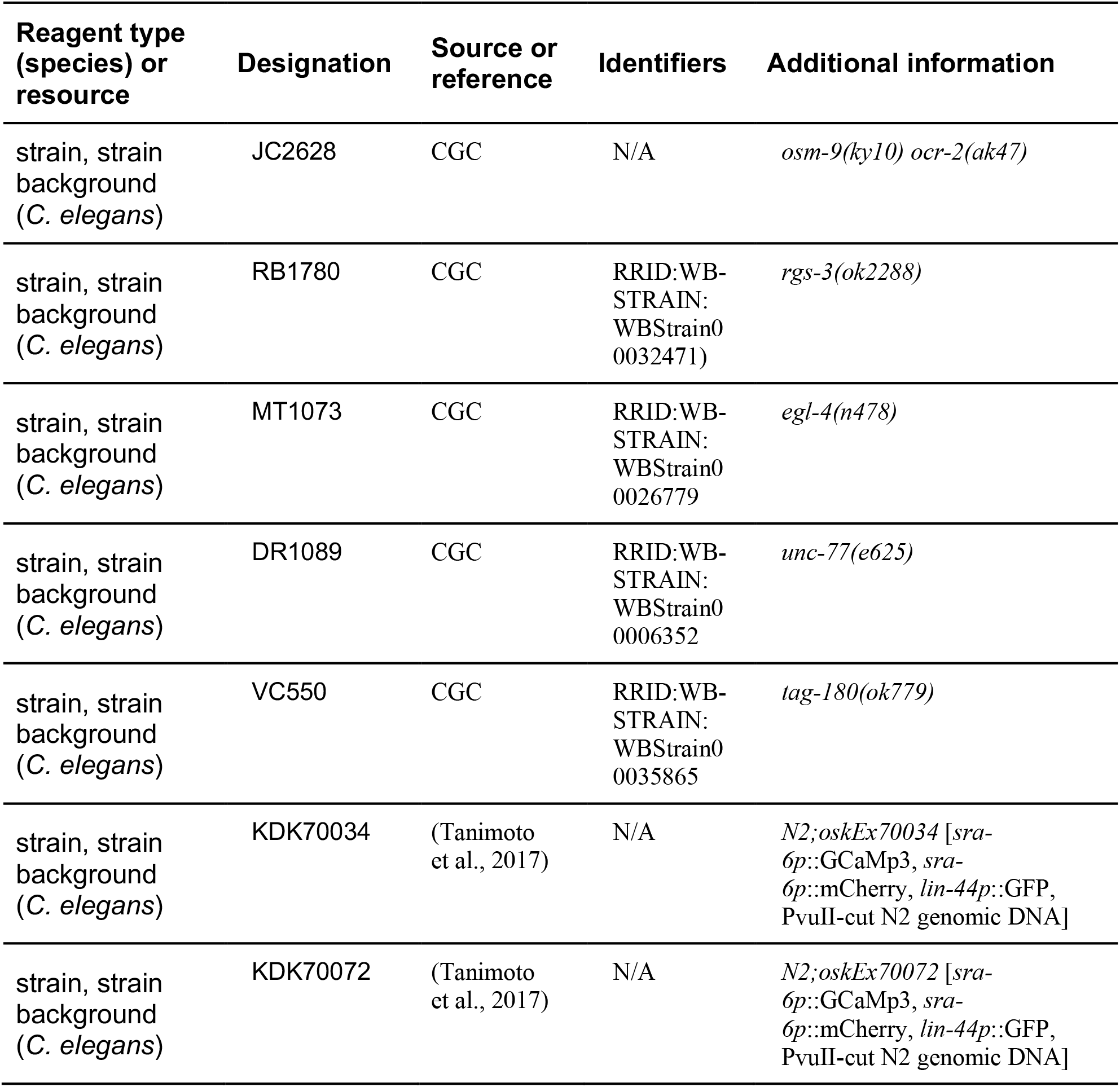
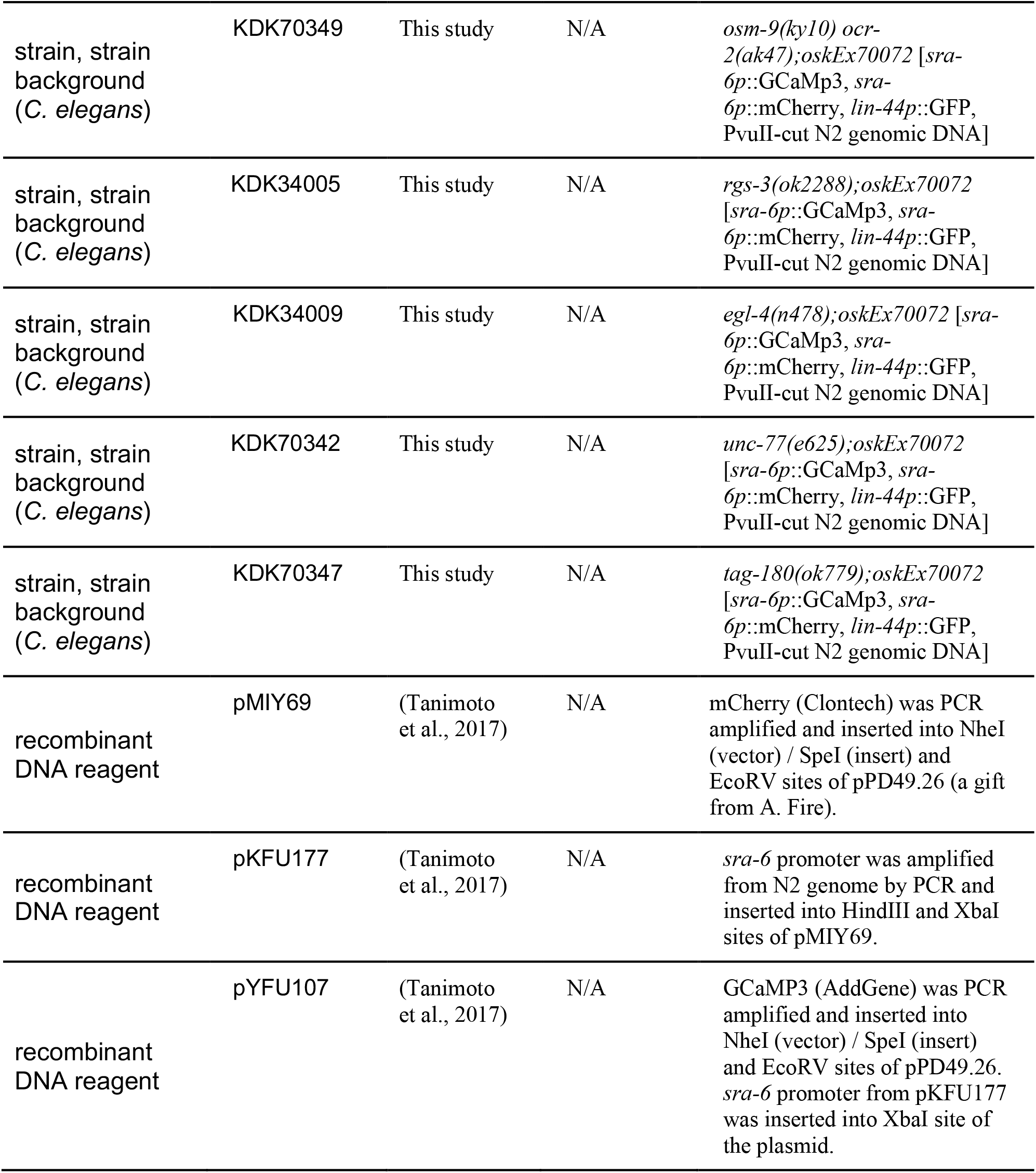

| Reagent type (species) or resource | Designation | Source or reference | Identifiers | Additional information |
| --- | --- | --- | --- | --- |
| strain, strain background (*C. elegans*) | JC2628 | CGC | N/A | *osm-9(ky10) ocr-2(ak47)* |
| strain, strain background (*C. elegans*) | RB1780 | CGC | RRID:WB-STRAIN: WBStrain0 0032471) | *rgs-3(ok2288)* |
| strain, strain background (*C. elegans*) | MT1073 | CGC | RRID:WB-STRAIN: WBStrain0 0026779 | *egl-4(n478)* |
| strain, strain background (*C. elegans*) | DR1089 | CGC | RRID:WB-STRAIN: WBStrain0 0006352 | *unc-77(e625)* |
| strain, strain background (*C. elegans*) | VC550 | CGC | RRID:WB-STRAIN: WBStrain0 0035865 | *tag-180(ok779)* |
| strain, strain background (*C. elegans*) | KDK70034 | (Tanimoto et al., 2017) | N/A | *N2;oskEx70034 [sra-6p::GCaMp3, sra-6p::mCherry, lin-44p::GFP, PvuII-cut N2 genomic DNA]* |
| strain, strain background (*C. elegans*) | KDK70072 | (Tanimoto et al., 2017) | N/A | *N2;oskEx70072 [sra-6p::GCaMp3, sra-6p::mCherry, lin-44p::GFP, PvuII-cut N2 genomic DNA]* |
| strain, strain background (*C. elegans*) | KDK70349 | This study | N/A | *osm-9(ky10) ocr-2(ak47);oskEx70072 [sra-6p::GCaMp3, sra-6p::mCherry, lin-44p::GFP, PvuII-cut N2 genomic DNA]* |
| strain, strain background (*C. elegans*) | KDK34005 | This study | N/A | *rgs-3(ok2288);oskEx70072 [sra-6p::GCaMp3, sra-6p::mCherry, lin-44p::GFP, PvuII-cut N2 genomic DNA]* |
| strain, strain background (*C. elegans*) | KDK34009 | This study | N/A | *egl-4(n478);oskEx70072 [sra-6p::GCaMp3, sra-6p::mCherry, lin-44p::GFP, PvuII-cut N2 genomic DNA]* |
| strain, strain background (*C. elegans*) | KDK70342 | This study | N/A | *unc-77(e625);oskEx70072 [sra-6p::GCaMp3, sra-6p::mCherry, lin-44p::GFP, PvuII-cut N2 genomic DNA]* |
| strain, strain background (*C. elegans*) | KDK70347 | This study | N/A | *tag-180(ok779);oskEx70072 [sra-6p::GCaMp3, sra-6p::mCherry, lin-44p::GFP, PvuII-cut N2 genomic DNA]* |
| recombinant DNA reagent | pMIY69 | (Tanimoto et al., 2017) | N/A | mCherry (Clontech) was PCR amplified and inserted into NheI (vector) / SpeI (insert) and EcoRV sites of pPD49.26 (a gift from A. Fire). |
| recombinant DNA reagent | pKFU177 | (Tanimoto et al., 2017) | N/A | *sra-6* promoter was amplified from N2 genome by PCR and inserted into HindIII and XbaI sites of pMIY69. |
| recombinant DNA reagent | pYFU107 | (Tanimoto et al., 2017) | N/A | GCaMP3 (AddGene) was PCR amplified and inserted into NheI (vector) / SpeI (insert) and EcoRV sites of pPD49.26. *sra-6* promoter from pKFU177 was inserted into XbaI site of the plasmid. |

### Strains

The techniques we used to culture and handle *C. elegans* were essentially as described previously (Brenner, 1974). *C. elegans* wild-type Bristol strain N2, JC2628 *osm-9(ky10) ocr-2(ak47)*, RB1780 *rgs-3(ok2288)*, MT1073 *egl-4(n478)*, DR1089 *unc-77(e625)*, and VC550 *tag-180(ok779)*, were obtained from the Caenorhabditis Genetics Center (University of Minnesota, USA). In all the behavioral and physiological experiments, we used young adult hermaphrodites.

### Calcium imaging by OSB2 system

Calcium imaging of ASH sensory neurons was performed as previously reported (Tanimoto et al., 2017; Tanimoto and Kimura, 2021). In brief, transgenic lines expressing GCaMP3 (Tian et al., 2009) and mCherry (Shaner et al., 2004) in ASH (20 ng/μl of *sra-6p*::GCaMP3, 20 ng/μl of *sra-6p*::mCherry, 10 ng/μl of *lin-44p*::GFP, 50 ng/μl of PvuII-cut N2 genomic DNA) were placed on NGM plates and observed under our original microscope system, OSB2. To measure the neural activity of multiple animals in a single observation, the transgenic animals were immobilized using levamisole, an agonist to the acetylcholine receptor (Lewis et al., 1980); ASH response is not affected by levamisole treatment (Tanimoto et al., 2017).

For odor stimulation, we delivered a mixture of 2-nonanone and air at a total of 8 mm/min and created a temporal changing gradient of odor concentration by changing the ratio of 2-nonanone to air. We measured this gradient using a custom-made semiconductor sensor before and after performing the calcium imaging experiments on each day (Tanimoto and Kimura, 2021).

We divided the fluorescent signals of GCaMP3 and mCherry through a dual-wavelength measurement optical system, w-view (Hamamatsu Photonics), and captured their fluorescence images using an EM-CCD camera (ImagEM, Hamamatsu Photonics) at a sampling rate of 10 Hz. After subtracting the background, we extracted the fluorescence intensity of the cell body using ImageJ (NIH) and used their ratio (GCaMP/mCherry) as the data.

### Conditioning of the animals

Calcium imaging of ASH was performed using OSB2 under the following three conditions. (1) Naive: the animals were cultivated on 6 cm NGM plates with OP50 provided as food, and were then washed briefly with NGM buffer and measured; (2) Preexp: the animals were preexposed to 0.6 μl of 15% 2-nonanone (diluted in ethanol) for 1 hour on a 6 cm NGM plate without food; and (3) Mock-treated: the animals were preexposed to ethanol in the same way as the preexp condition. The mock-treated condition is a control that shows that starvation itself does not affect the response to 2-nonanone.

### Data analysis and statistical analysis

In the calcium imaging experiments, data were obtained two to three days in each condition. Experimental conditions, such as strains, odor stimuli and conditioning of the animals, were randomly set for each day. After the data acquisition, animals with too weak intensity of basal mCherry or GCaMP3 fluorescence for tracking were excluded. To exclude noise, frames in which the GCaMP/mCherry ratio was in the top 1% and bottom 1% of the total were removed. Statistical analyses were performed using the Kruskal-Wallis test with a *post-hoc* Steel-Dwass test using R (The R Project). In the figure, * indicates *p* < 0.05, ** indicates *p* < 0.01, and *** indicates *p* < 0.001. Scatter plots represent median ± quartiles and other graphs represent the mean ± standard error. We chose the sample size based on a large scale behavioral analysis of *C. elegans* (Yemini et al., 2013).

### Mathematical modeling of AWB-like responses by leaky integration

Mathematical modeling of AWB-like responses was described in our previous work (Tanimoto et al., 2017). In brief, the activity of AWB neurons, which responds to decreases in 2-nonanone concentration, is well fitted with the following leaky integrator equation:

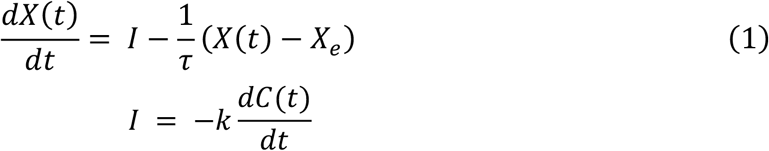

where *X(t)* is the measured fluorescence signal of the neuron (GCaMP/mCherry), *I* is the input, *X*_*e*_ is the basal calcium level in the steady state, *k* and 1/*τ* are the parameters, and *C(t)* is the measured odor concentration. The model parameters *k*, 1/*τ*, and *X*_*e*_ were determined by the least-squares method (Excel solver). This Eq. (1) indicates that the neuronal response *X(t)* increases according to the input (negative *dC(t)/dt*) and reduces (“leaks”) according to the *X(t)* itself and returns to the steady state *X*_*e*_. For example, when an animal experiences a constant decrease in odor concentration (*e*.*g*., Figure 1—figure supplement 2C), the constant decrease is transformed to a constant depolarization, and it causes constant calcium influx via L-type VGCC EGL-19, and the calcium concentration reduces according to its total amount possibly via leak channel(s) (Tanimoto et al., 2017). Since AWB neurons respond to decreases in odor concentration, zero is substituted to *dC(t)/dt* when *dC(t)/dt* is positive in Eq. (1). This AWB activity was essentially abolished by treatment of animals with Nemadipine-A (NemA), the antagonist of *C. elegans* L-type VGCC (Kwok et al., 2006), indicating that the AWB activity is mainly mediated by the L-type VGCC homolog EGL-19.

In NemA-treated animals, ASH neurons are rapidly activated at the beginning of the odor-increasing phase but soon inactivated gradually even though the odor concentration kept increasing (red dotted line in Figure 1—figure supplement 2D). When we assumed that the NemA-treatment suppressed AWB-like leaky integration activity (yellow dotted line in Figure 1—figure supplement 2D), the sum of the remaining and the suppressed activity (blue solid line in Figure 1—figure supplement 2D) nicely resembled the measured ASH activity (black solid line in Figure 1—figure supplement 2D). We call the transient and gradual activities the fast and slow components, respectively (Tanimoto et al., 2017).

### The first and second differential model of ASH responses

We extended the previous model Eq. (1) by introducing the second-order time differential of input *C* representing the fast component:

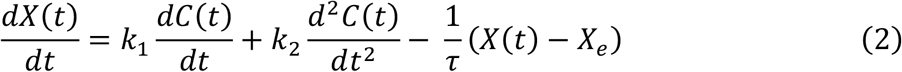

where *X(t)* is the measured ASH signal, *X*_*e*_ is the basal calcium level, and *k*_*1*_, *k*_*2*_ and 1/*τ* are parameters for the first- and second-order time differential and leaky part, respectively. The parameters *k*_*1*_, *k*_*2*_, 1/*τ*, and *X*_*e*_ were determined by the least-squares method for each odor stimulus. *dC(t)/dt* and *d*^*2*^*C(t)/dt*^*2*^ were calculated using the central difference scheme. Since ASH neurons respond to increases in odor concentration (Tanimoto et al., 2017), zero is substituted to *dC(t)/dt* or *d*^*2*^*C(t)/dt*^*2*^ when *dC(t)/dt* or *d*^*2*^*C(t)/dt*^*2*^ is negative in Eq. (2).

The first and second differential model Eq. (2) is further extended on the terms of the second-order time-differential and *X*_*e*_:

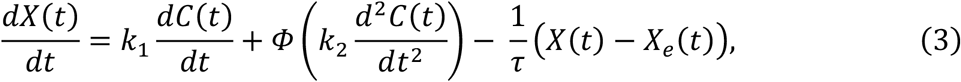

where *Φ(x)* is the saturation function as the following equation:

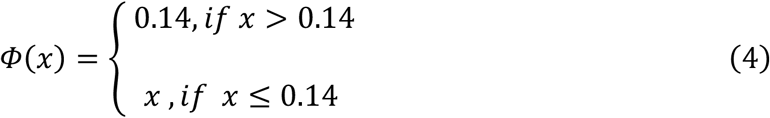

This saturation function is introduced to prevent the parameters from becoming too large when the increase in odor concentration approaches zero. 0.14 was arbitrary determined. *X*_*e*_*(t)* changes over time as follows:

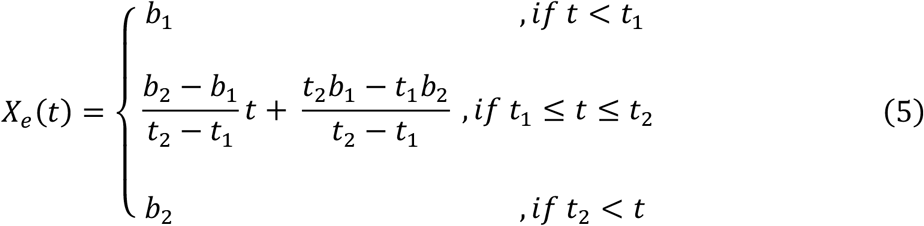

where *t*_*1*_ is the time at the start of the first odor increase phase, and *t*_*2*_ is the time at the end of the first odor increase phase. The parameters *b*_*1*_ and *b*_*2*_ were determined by the least-squares method with the actual ASH response during 30 seconds until 10 seconds before the start of the first odor increase phase (*t*_*1*_) and 1 minute after the first odor stimulus (*t*_*2*_), respectively. Eq. (5) was introduced because the basal calcium level changed after the first odor stimulus.

In terms of parameters in Eq. (3), we plotted the relationships among parameters (*k*_*1*_, *k*_*2*_, and 1/*τ*) and various aspects of odor inputs (average, maximum, or accumulation of *C, dC/dt*, and *d*^*2*^*C/dt*^*2*^) and fitted the relationships to the logarithmic function (y = *f* ln(x) + *g*) of the odor inputs (Excel solver) (Figure 3A and Figure 3—figure supplement 2); fitting of *k*_*1*_ and 1/*τ* to linear functions (y = *h* x + *i*) of the odor inputs was worsened (Table 3). We chose the aspect of odor input with the smallest or the second smallest residual sum of squares, and the logarithmic function of our chosen aspect for *k*_*1*_ and 1/*τ* were similar between naive, mock-treated and preexposed conditions. Thus we further optimized *f* and *g* manually such that they were the same for all three conditions.

**Table 3.** Comparison of BIC for linear and logarithmic approximation of stimulus-dependent parameters.

| Odor | #1 | #2 |  |  |
| --- | --- | --- | --- | --- |
| Condition | Naive | Naive | Mock-treated | Preexp |
| Linear approximation | -539.2 | -899.6 | -1009 | -682.0 |
| Logarithmic approximation | **-778.4** | **-1135** | **-1116** | **-966.9** |

The conditions with the smallest BIC values under each condition are indicated in bold. In all the conditions, the model with a
logarithmic approximation has a better BIC value than the model with a linear approximation.

As a lower threshold for 1/*τ*, the following function is introduced:

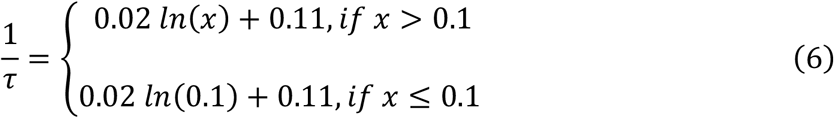

where *x* denotes *d*^*2*^*C(t)/dt*^*2*^ (Figure 3A). This function is introduced to prevent 1/*τ* from becoming too small when the increase in odor concentration approaches zero. 0.1 was arbitrary determined.

### Evaluation of the mathematical model

The Bayesian information criterion (BIC) (Schwarz, 1978) was used to assess the fit of the data using a mathematical model. The smaller the BIC value, the better is the model fit. In BIC, the goodness of fit for the model, including a penalty term to prevent overfitting, is given by the following equation:

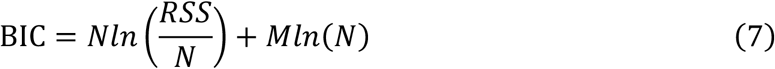

where *N* is the number of frames used for the fitting, *RSS* is the sum of the squared residuals between the model and the actual response, and *M* is the number of free parameters used in the model. *M*=2 for the original simple time-differential model (Figure 2—figure supplement 1) because of *k* and *X*_*e*_, *M*=3 for the model with the leaky integration of only first- or second-order time-differential (Figure 2—figure supplement 2 and 3) because of *k*_*1*_ or *k*_*2*_ and 1/*τ* and *X*_*e*_, *M*=4 for the first and second differential model (Figure 2B) because of *k*_*1*_, *k*_*2*_, 1/*τ* and *X*_*e*_, and *M*=12 for the first and second differential model with stimulus-dependent parameters (Figure 3B) because there were two parameters each in the stimulus-dependent parameters *k*_*1*_, *k*_*2*_, and 1/*τ*, four parameters in *X*_*e*_*(t)*, one parameter in *Φ(x)*, and one parameter in lower threshold for 1/*τ*.

### Behavioral analysis

Trajectories obtained by a high-resolution USB camera (DMK72AUC02; The Imaging Source, United States) during 2-nonanone avoidance behavior were clustered using the previously reported STEFTER method (Yamazaki et al., 2019). In brief, clustering was performed using variances of temporal changes in bearing, and the cluster with the smallest variances of temporal changes in bearing was classified as “run”, while the other clusters were classified as “pirouette” categories. This classification of “run” and “pirouette” was more than 90% consistent with the previous one based on the durations of straight migration (Yamazoe-Umemoto et al., 2015). Then, the change in the odor concentration sensed during the run was calculated according to an odor gradient model based on the measured odor concentrations (Yamazoe-Umemoto et al., 2015). The behavioral data used in this study have already been analyzed and published previously (Yamazaki et al., 2019).

## ACKNOWLEDGEMENTS

We thank Yuki Tsukada, Yuishi Iwasaki, Sakiko Shiga, Masahiro Ueda, and the other Kimura laboratory members for their valuable advice, comments and technical assistance for this study. Nematode strains were provided by the Caenorhabditis Genetics Center (funded by the NIH Office of Research Infrastructure Programs P40 OD010440).

