## Supplementary Figures for "Neural mechanism of experience-dependent sensory gain control in *C. elegans*"

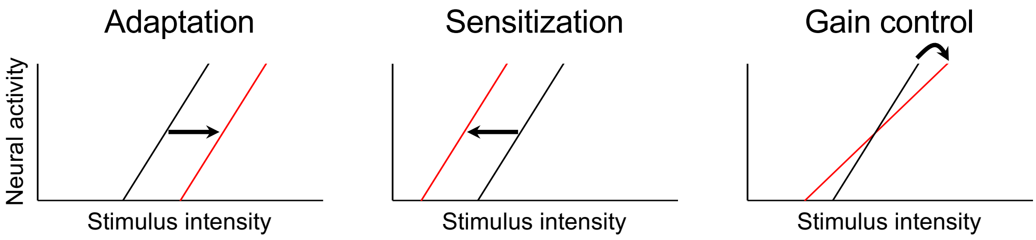


**Figure 1—figure supplement 1:** Schematic drawing of sensory modulations. Adaptation shifts the stimulus–response relationship to the right and sensitization shifts it to the left. Gain control is a change of its slope.

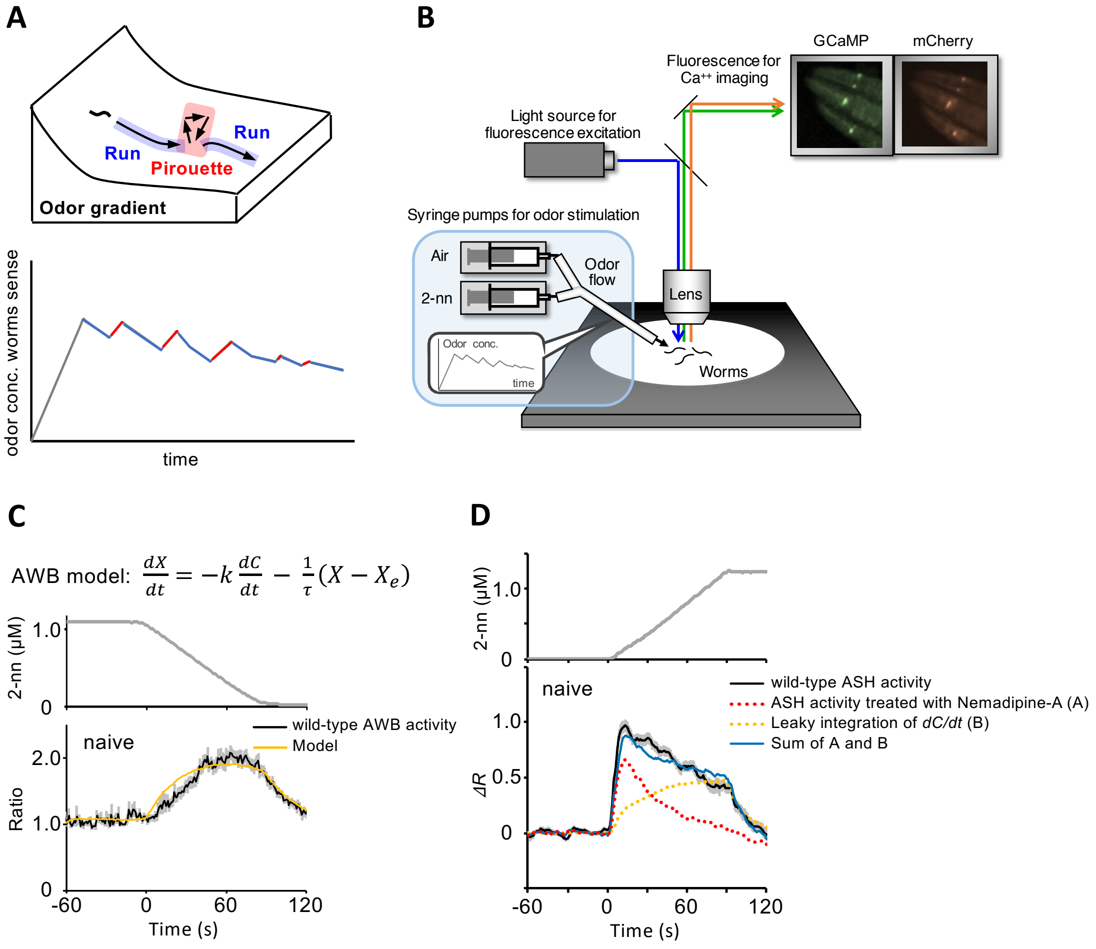


**Figure 1—figure supplement 2:** Behavioral and neuronal responses of *C. elegans* to the repulsive odor 2-nonanone. **A,** Odor concentration changes during avoidance behavior. (Top) Schematic drawing of a 2-nonanone gradient and the animals' behavioral states (runs and a pirouette) during the avoidance behavior. (Bottom) An example of odor concentration changes sensed by an animal during avoidance assay and the behavioral state during these changes. Runs (blue) and pirouettes (red) are mainly associated with a decrease and increase in odor concentration. In the initial phase of the assay (gray), the odor concentration always increases because the odor rapidly evaporates from its source. **B,** Schematic drawing of calcium imaging by our original microscope system OSB2. Worms immobilized with the acetylcholine receptor antagonist levamisole were exposed to an odor flow, and the neural activities were monitored with dual color imaging. **C,** AWB response to a constant gradual decrease in 2-nonanone concentration. When a constant odor decrease (upper panel) is presented,

AWB neurons in naive wild-type animals exhibits a gradually increasing response (black solid line in lower panel), which can be approximated by the leaky integration equation (top and yellow line in lower panel). The details of leaky integration equation are described in Materials and Methods. **D,** ASH response to a constant gradual increase in 2-nonanone concentration. When a simple gradual odor increase (upper panel; odor gradient #1) is presented, ASH neurons in naive wild-type animals exhibit a fast and relatively constant response (black solid line in lower panel). In contrast, when wild-type animals are treated with Nemadipine-A, a specific antagonist for L-type VGCC EGL-19, ASH exhibits a fast and transient response (red dashed line in lower panel). Interestingly, when the transient response and the calculated leaky integration of time-differential of odor concentration mediated by the EGL-19 (yellow dashed line) are added (blue dashed line), the added result nicely reproduces the actual ASH response. For panel D, instead of the ratio (GCaMP/mCherry), $\Delta R$ (ratio - baseline) was used because in this panel, two different groups of wild-type ASH responses (untreated versus NemA-treated) were compared. Panels A–D are reproduced from the previous studies with some modifications (Tanimoto et al., 2017; Yamazaki et al., 2019).

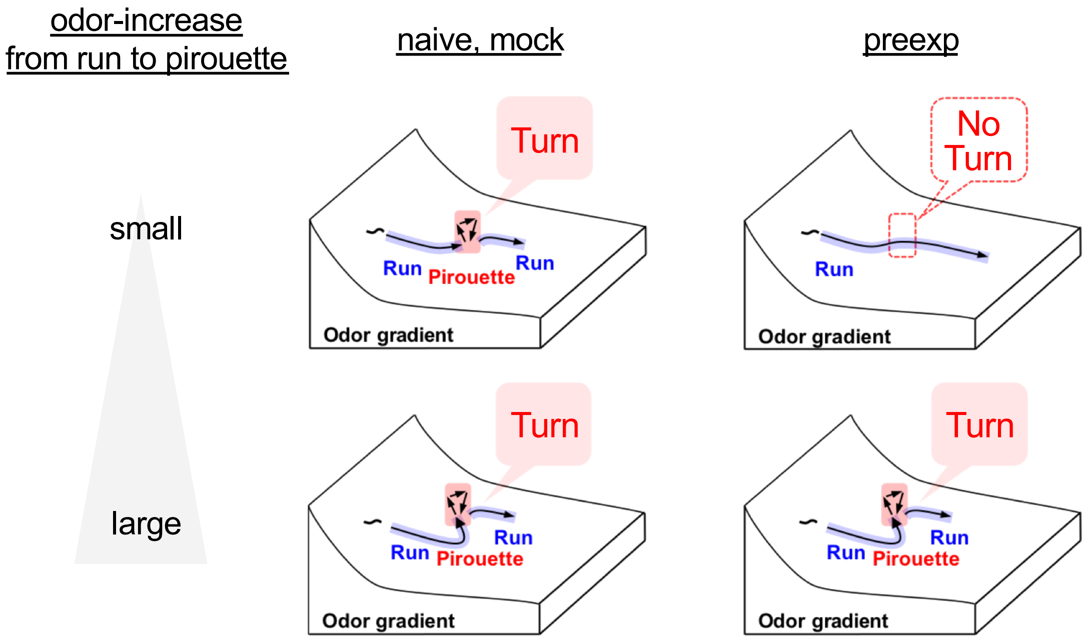


**Figure 1—figure supplement 3:** Cartoons of experience-dependent modulation of the odor avoidance behavior. An efficient repulsive behavior can be accomplished by only suppressing the response to a slight odor increase. A part of this figure has been published previously (Yamazaki et al., 2019).

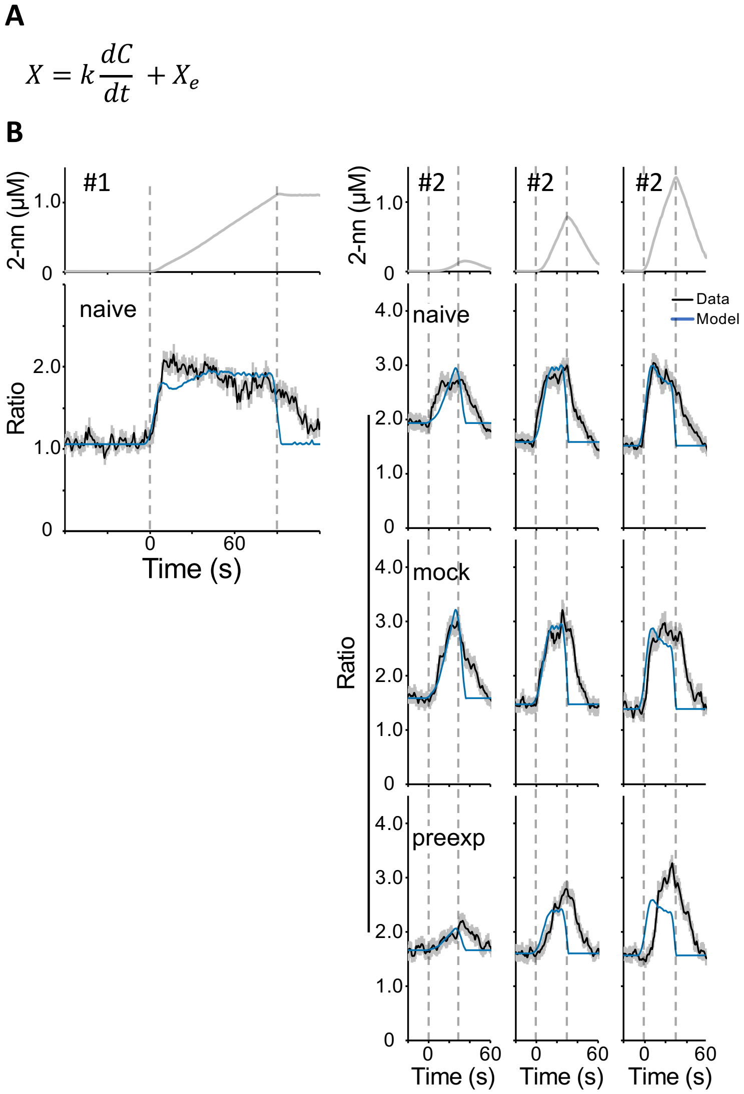


**Figure 2—figure supplement 1:** Independent fitting of the wild-type ASH response to the odor stimuli with the original simple time-differential model. **A,** Equation of the model. **B,** The fitting results of the ASH response to the odor gradient #1 (left) and to each stimulus of the odor gradient #2 (right). The black line indicates the actual ASH response and the blue line is the model. The black dotted line indicates the onset and end of the odor increase phase.

**A**

$$\frac{dX}{dt} = k_1 \frac{dC}{dt} - \frac{1}{\tau}(X - X_e)$$

**B**

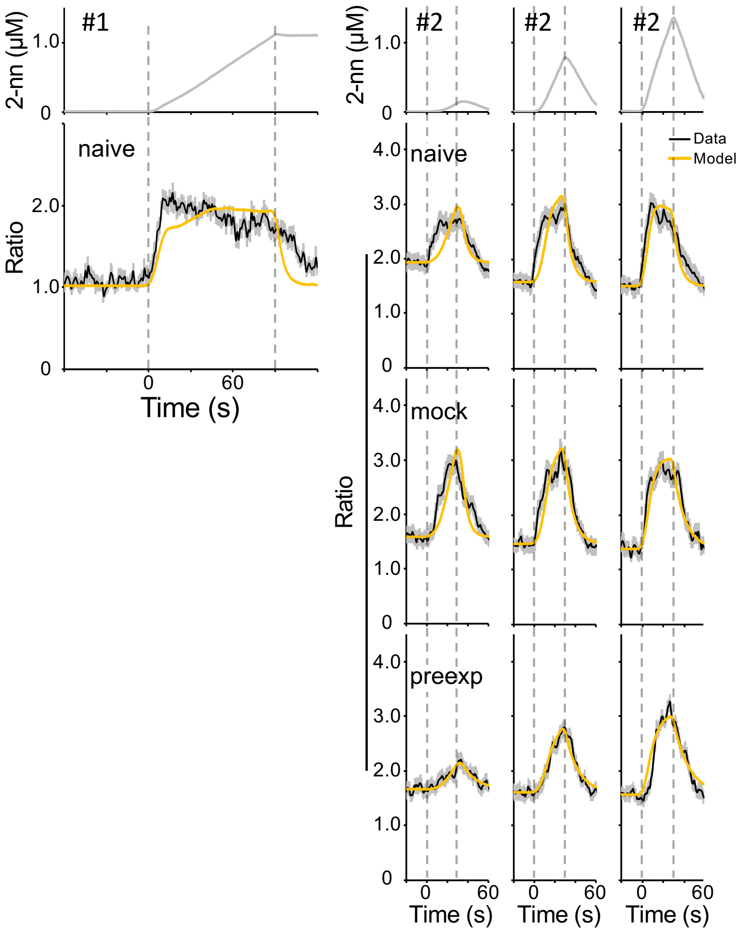


**Figure 2—figure supplement 2:** Independent fitting of the wild-type ASH response to the odor stimuli with the leaky integration of first-order time-differential model. **A,** Equation of the model. **B,** The fitting results of the ASH response to the odor gradient #1 (left) and to each stimulus of the odor gradient #2 (right). The black line indicates the actual ASH response and the yellow line is the model. The black dotted line indicates the onset and end of the odor increase phase.

**A**

$$\frac{dX}{dt} = k_2 \frac{d^2C}{dt^2} - \frac{1}{\tau}(X - X_e)$$

**B**

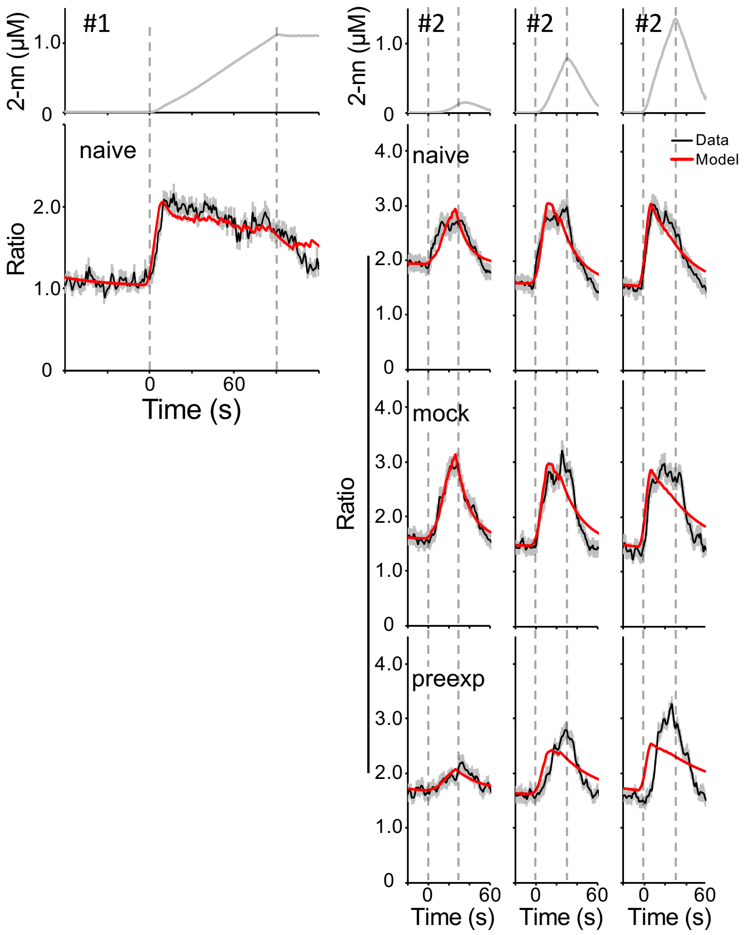


**Figure 2—figure supplement 3.** Independent fitting of the wild-type ASH response to the odor stimuli with the leaky integration of second-order time-differential model. **A,** Equation of the model. **B,** The fitting results of the ASH response to the odor gradient #1 (left) and to each stimulus of the odor gradient #2 (right). The black line indicates the actual ASH response and the red line is the model. The black dotted line indicates the onset and end of the odor increase phase.

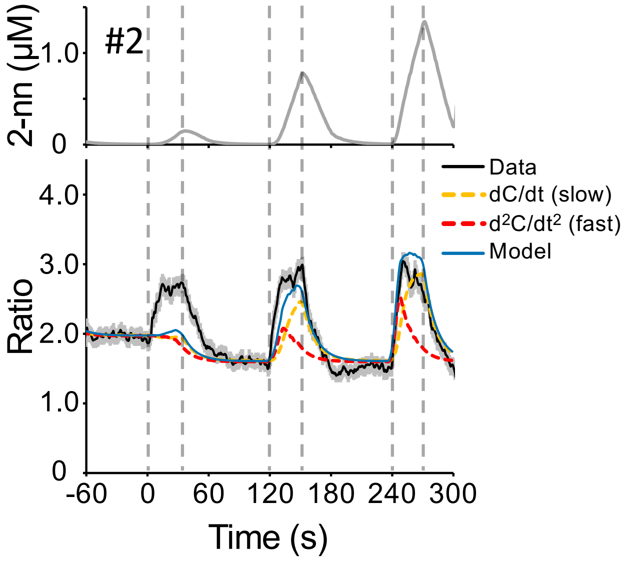


**Figure 3—figure supplement 1:** Fitting of the model with constant parameters for whole naive ASH response to the odor gradient #2. The model was fitted through all odor stimuli based on the least-squares method. The responses to 20 nM/s and 40 nM/s increases were well reproduced while the one to 3 nM/s increase was not.

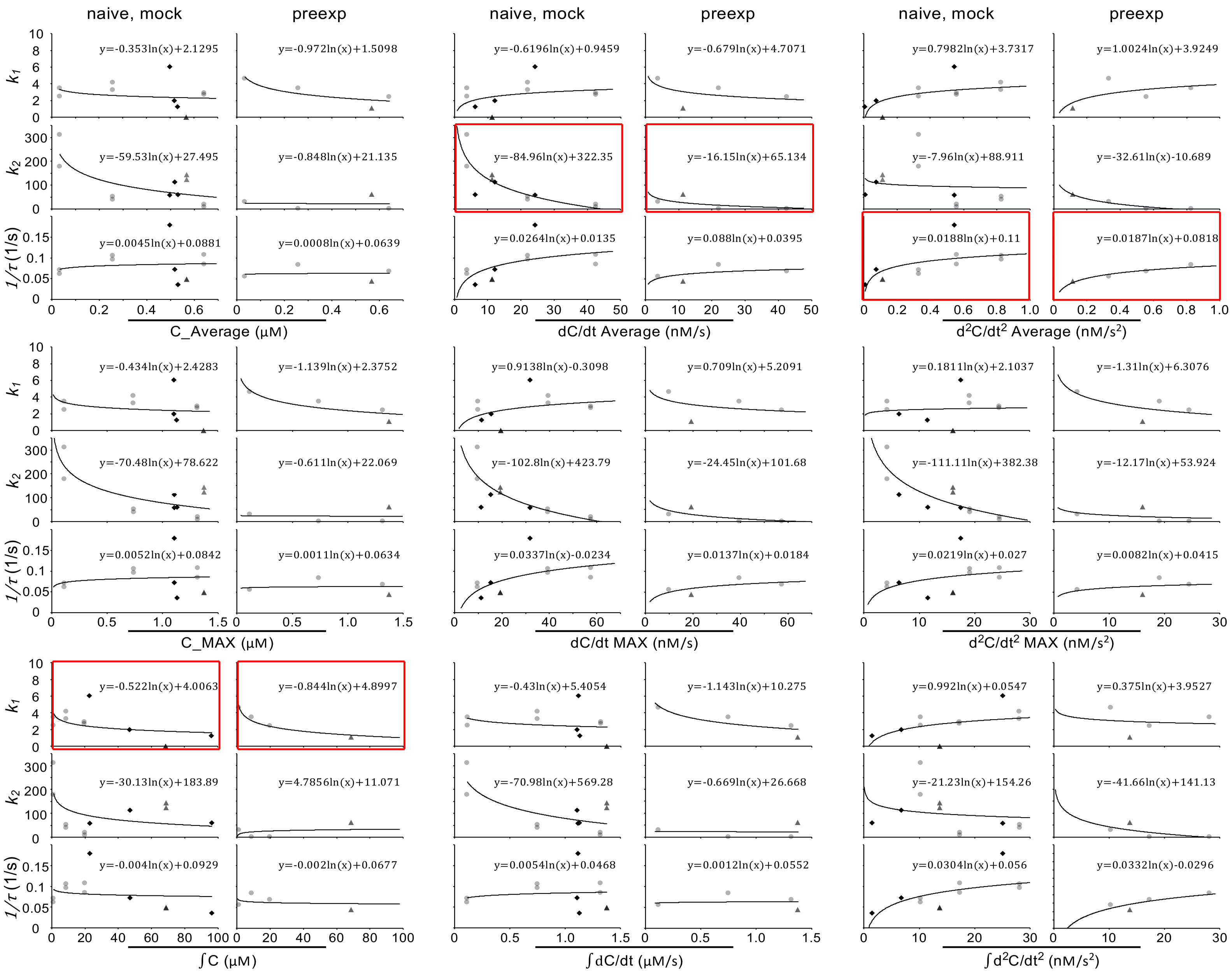
naive, mock
preexp
naive, mock
preexp
naive, mock
preexp
$k_1$
$k_2$
$1/\tau$ (1/s)
y=-0.353ln(x)+2.1295
y=-0.972ln(x)+1.5098
y=-0.6196ln(x)+0.9459
y=-0.679ln(x)+4.7071
y=0.7982ln(x)+3.7317
y=1.0024ln(x)+3.9249
y=-59.53ln(x)+27.495
y=-0.848ln(x)+21.135
y=-84.96ln(x)+322.35
y=-16.15ln(x)+65.134
y=-7.96ln(x)+88.911
y=-32.61ln(x)-10.689
y=0.0045ln(x)+0.0881
y=0.0008ln(x)+0.0639
y=0.0264ln(x)+0.0135
y=0.088ln(x)+0.0395
y=0.0188ln(x)+0.11
y=0.0187ln(x)+0.0818
C_Average (μM)
dC/dt Average (nM/s)
d$^2$C/dt$^2$ Average (nM/s$^2$)
y=-0.434ln(x)+2.4283
y=-1.139ln(x)+2.3752
y=0.9138ln(x)-0.3098
y=0.709ln(x)+5.2091
y=0.1811ln(x)+2.1037
y=-1.31ln(x)+6.3076
y=-70.48ln(x)+78.622
y=-0.611ln(x)+22.069
y=-102.8ln(x)+423.79
y=-24.45ln(x)+101.68
y=-111.11ln(x)+382.38
y=-12.17ln(x)+53.924
y=0.0052ln(x)+0.0842
y=0.0011ln(x)+0.0634
y=0.0337ln(x)-0.0234
y=0.0137ln(x)+0.0184
y=0.0219ln(x)+0.027
y=0.0082ln(x)+0.0415
C_MAX (μM)
dC/dt MAX (nM/s)
d$^2$C/dt$^2$ MAX (nM/s$^2$)
y=-0.522ln(x)+4.0063
y=-0.844ln(x)+4.8997
y=-0.43ln(x)+5.4054
y=-1.143ln(x)+10.275
y=0.992ln(x)+0.0547
y=0.375ln(x)+3.9527
y=-30.13ln(x)+183.89
y=4.7856ln(x)+11.071
y=-70.98ln(x)+569.28
y=-0.669ln(x)+26.668
y=-21.23ln(x)+154.26
y=-41.66ln(x)+141.13
y=-0.004ln(x)+0.0929
y=-0.002ln(x)+0.0677
y=0.0054ln(x)+0.0468
y=0.0012ln(x)+0.0552
y=0.0304ln(x)+0.056
y=0.0332ln(x)-0.0296
∫C (μM)
∫dC/dt (μM/s)
∫d$^2$C/dt$^2$ (nM/s$^2$)

**Figure 3—figure supplement 2:** Scatter plots of the relationship between each aspect of stimuli and parameter. We plotted the relationship between the parameters that reproduced the ASH response to each odor increase and their inputs (average, maximum and integrated value of odor concentration, first-order time-differential of odor concentration, and second-order time-differential of odor concentration). Black rhombuses, light gray circles, and dark gray triangles represent ASH responses to odor gradient #1, #2, and #3, respectively. The relationships finally chosen (Figure 3A) are indicated by red rectangles.
